# Reproducibility of the evaluation of genetic variant pathogenicity based on the animal variant classification guidelines

**DOI:** 10.1101/2025.11.12.687631

**Authors:** Iris Casselman, Carlotta Ferrari, Marie Abitbol, Danika Bannasch, Jerold Bell, Caroline Dufaure de Citres, Carrie J. Finno, Jessica J. Hayward, Jens Häggström, Jason T. Huff, Tosso Leeb, Ingrid Ljungvall, Maria Longeri, Leslie A. Lyons, Marcela Martinez, Cathryn Mellersh, Frank W. Nicholas, Åsa Ohlsson, Pascale Smets, Maria G. Strillacci, Imke Tammen, Frank G. van Steenbeek, Bart J.G. Broeckx

## Abstract

Until recently, due to the absence of standardized guidelines tailored for veterinary use, the evaluation of genetic variant pathogenicity for single-gene diseases was based on a personal interpretation of the presented evidence, which has led to ambiguous interpretation. With the publication of the animal variant classification guidelines (AVCG), a more objective approach became available. Variants are evaluated based on twenty-three criteria and labeled as pathogenic, likely pathogenic, variant of uncertain significance, likely benign or benign. While the accuracy was thoroughly tested in the original publication, the reproducibility of the various steps involved was only briefly checked, which is why the current study was performed. Each variant from a set of 150 published likely causal variants for single-gene diseases from three species (dog, cat, horse) was independently and blindly assessed by three different reviewers, each applying the same AVCG. An overall agreement of 93% for decisions on the scope, i.e. whether they fit the inclusion criteria to allow evaluation with AVCG, was found. More importantly, the reproducibility was 65% for the pathogenicity classification and this increased to 83% clinically relevant agreement. While a direct comparison of the reproducibility with human literature is not possible for the scope, the reproducibility on pathogenicity classification is in line with reports using the human American College for Medical Genetics and Genomics and Association for Molecular Pathology guidelines for human variants. Overall, based on the current study, the reproducibility of the guidelines in veterinary species is within current expectations.

## Introduction

Only a subset of all the genetic variation naturally present in populations causes diseases (Amendola et al., 2020). While technologies have simplified the discovery of genetic variants, discriminating the disease-causing variants remains difficult for a combination of reasons (Boeykens et al., 2023). Until recently, animal geneticists mostly relied on their own expertise to decide when a pathogenicity threshold had been met (Boeykens et al., 2024b). Inherently a subjective method, differences between evaluators and laboratories are likely and have been reported in human and veterinary genetics (Amendola et al., 2016; Boeykens et al., 2024a; Yorczyk et al., 2015). In human genetics, the American College for Medical Genetics and Genomics and Association for Molecular Pathology (ACMG/AMP) guidelines were developed to resolve this challenge and provide guidance on when the pathogenicity support was sufficient to use a variant in clinical decision-making (Richards et al., 2015). Recently, a first version of the Animal Variant Classification Guidelines (AVCG) tailored for veterinary use was published (Boeykens et al., 2024a) and since then, these guidelines have been used several times to classify new disease-causing variants (Abitbol et al., 2025; Boeykens et al., 2025, 2024a; Kaczmarska et al., 2025; Stee et al., 2025). Briefly, this system is based on 23 criteria, with variable weights (“very strong”, “strong”, “moderate”, “supportive”, see Suppl. File 1) (Boeykens et al., 2024a). Based on a decision tree, a certain combination of criteria leads to one of five labels, which are the same as the ACMG/AMP labels (pathogenic (P), likely pathogenic (LP), variant of uncertain significance (VUS), likely benign (LB) and benign (B)) (Suppl. File 2) (Boeykens et al., 2024a; Richards et al., 2015). These labels are important as they influence clinical decision-making, with P and LP being actionable and the other three not actionable (Suppl. File 3) (Amendola et al., 2020, 2016; Boeykens et al., 2024a; Rehm et al., 2015; Richards et al., 2015; Rius et al., 2025). As with every classification system, these guidelines need to be further evaluated to quantify various performance characteristics. An important characteristic is the interobserver agreement, or reproducibility, which quantifies the consistency of classification among different evaluators. A high reproducibility should not be taken for granted, as numerous studies prove. Reproducibility can be influenced by many factors, such as the classification system, the phenotype studied, and educational attainment (Amendola et al., 2016; Bertal et al., 2018; Boeykens et al., 2024b; Bogaerts et al., 2018; De Vos et al., 2025; Rehm et al., 2015; Verhoeven et al., 2007; Yorczyk et al., 2015).

While the original study detailing the AVCG evaluated reproducibility to some extent, the dataset contained only 17 variants from one species, which was not sufficient for an in-depth evaluation. Performed under the umbrella of the Animal Genetic Testing Standardization (AGTS) Committee of the International Society for Animal Genetics (ISAG), the aim of the current study is to evaluate the reproducibility of AVCG using a large dataset across several species and to identify factors that might improve the classification process.

## Materials and methods

### Reviewers

All authors involved in the original AVCG publication were asked whether they wanted to continue (Boeykens et al., 2024a). An invitation to participate was sent via email to additional members of the ISAG community. To allow a comparison with human literature, only experienced reviewers, defined as animal geneticists with at least 10 years of professional experience in the field of veterinary genetics, were selected to review variants. Each reviewer had to specify for which species they were comfortable classifying variants. Reviewers who indicated one species were assigned that species, while reviewers who indicated more than one species, were assigned to their preferred one or two species in such a way that there were three reviewers per variant. In total, 15 reviewers performed the actual classifications. One additional spare reviewer (reviewer #16) was available to classify variants for which there was a conflict of interest (COI), as detailed below.

### Variant source

Variant information was downloaded from the Online Mendelian Inheritance in Animals (OMIA) database website (Nicholas et al., 2025; Tammen et al., 2024), subsection “*All known likely causal variants for single-gene diseases*”, alphabetically sorted on gene name, on the 10^th^ of January, 2025 (dog) and on the 14^th^ of January, 2025 (horse and cat), until a sufficient number of variants was available, i.e. 30 variants per reviewer from their preferred species. Variants were randomly allocated to reviewers.

### Evaluation of variants

The evaluation process was divided into an administrative part, a check whether the variants fell in the scope, and the actual evaluation of the variant (Fig. 1). To aid in the process, every evaluator received a manual containing the criteria (Suppl. File 1), the clarifications and the decision table (Suppl. File 2), as detailed in Boeykens et al., 2024a, as well as an Excel file where the evaluation had to be filled in.

**Fig. 1.**
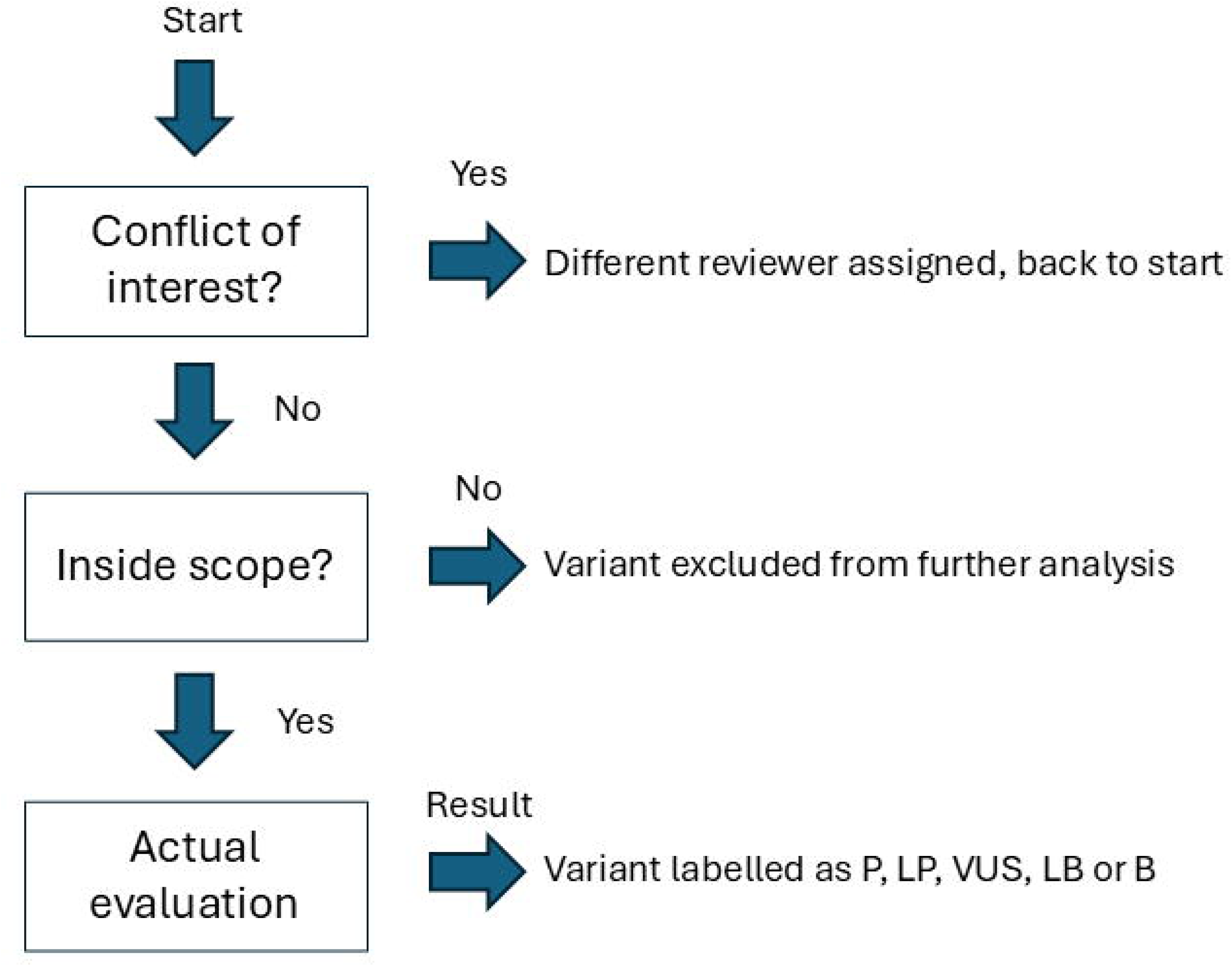
Overview of the evaluation process, which was divided into an administrative part (checking for a conflict of interest), check of the scope and actual evaluation of an individual variant. Abbreviations: P = pathogenic; LP = likely pathogenic; VUS = variant of uncertain significance; LB = likely benign; B = benign.

In the administrative part, a reviewer had to declare for each variant whether there was a COI, defined as “*being mentioned in the acknowledgements and/or author list*”. If a COI was identified, the spare reviewer (reviewer #16) was assigned, and the procedure was restarted. Next, it was checked whether the variant fell in the scope of AVCG, which was defined as “*The scope are single-gene disorders. These are disorders characterized by variants in a single gene with a high impact on disease risk, i.e. the impact of one variant is sufficient to cause disease (but does not have to cause disease in every individual due to e.g. reduced penetrance)”*. If a variant was determined to be in the scope, the actual evaluation was started. This involved the aforementioned evaluation of 23 criteria and assignment of a pathogenicity label based on a decision tree (ordered categories: P/LP/VUS/LB/B) (Boeykens et al., 2024a). The criteria, decision tree and explanation on how to interpret the labels, are available in Suppl. File 1-3, respectively.

Each step (i.e. COI, scope, criteria, label) was done for each variant by three reviewers. If a variant was considered outside the scope by ≥1 of the reviewers, it was excluded from further analyses. Within each species, variants were assigned randomly to the different reviewers and the classification process was done blinded (i.e. every reviewer classified every variant independently and was unaware of who of the other reviewers were evaluating that same variant, as well as what their evaluation was) to avoid e.g. influences caused by power relations (Boeykens et al., 2024a). The entire process is outlined in Fig. 1.

### Analysis

All analyses were performed using custom scripts in R version 4.3.2. The analysis was divided into a comparison of decisions on the scope, the individual criteria, and the final pathogenicity label (before and after diverging label correction, when applicable (see below)). To evaluate the agreement, the answers were compared pairwise. Practically, if a label/criterion/decision on the scope was the same for two reviewers, this was counted as an agreement. If the answer was not the same, it was counted as a disagreement. This was repeated for all three pairwise combinations (i.e. result of reviewer one vs reviewer two, reviewer two vs reviewer three, and reviewer one vs reviewer three) for a label/criterion/decision on the scope for every variant, and resulted in a range from 0 to 3 agreements per variant for a label/criterion/decision on the scope. By counting the number of agreements and dividing this by the theoretical maximum number of agreements (i.e. the number of comparisons performed, which was three times the number of variants), an agreement score was calculated. Specifically for the labels, more detailed evaluations were also performed, in agreement with human standards, i.e. classifications that might lead to medical management differences (i.e. P/LP vs B/LB/VUS) and classification differences associated with the confidence of the call (P vs LP; B vs LB; VUS vs LB/B) were further distinguished. This was done by grouping the labels as mentioned in the brackets and quantifying pairwise agreement (Amendola et al., 2020, 2016; Frone et al., 2021; Harrison et al., 2017; Maxwell et al., 2016; Sorelle et al., 2020). The agreement based on the former grouping is called “clinically relevant agreement”. In addition, the level of disagreement was also calculated, with a one-level disagreement defined as a one-category difference between the ordered pathogenicity levels (e.g. P vs LP) (Amendola et al., 2016). Using this system, the maximum difference is four levels.

Throughout the analysis, it became clear that tabulation errors had occurred sometimes. To evaluate whether the label of a reviewer corresponded with the combination of fulfilled criteria, an automated checking algorithm that assigned the label exactly based on the decision table, was developed. If a difference was identified, the reviewer was contacted and asked whether this was deliberate or a tabulation error and whether a correction had to be made or whether the original label should stay.

## Results

One hundred and fifty variants were included and assigned across 15 experienced geneticists, leading to a total of 450 evaluations. For six of these evaluations, a COI was reported, and the evaluation was assigned to a different reviewer (i.e. reviewer #16). There was no COI for any of these six variants for reviewer #16. Fifteen times (3%, 15/450), an individual variant was considered outside the scope by one reviewer, leading to fifteen variants being excluded (Suppl. File 4). The pairwise interobserver agreement on the scope was 93% (420/450 identical evaluations on the scope).

After exclusion of the variants outside the scope, 135 variants/ 405 classifications remained. Of these, 288 were P, 72 LP, 40 VUS, 1 LB and 4 were deliberately not assigned a label. Out of 135 variants, classification was unanimous for 73 variants (54%), while 45 (33%) and 17 (13%) variants received two or three different labels, respectively. Overall, this leads to a pairwise agreement of 65% (264/405). Of the 141 of 405 (35%) disagreements, 50% (71/141) were linked to the confidence of calls (P vs LP: 70/71; VUS vs LB: 1/71), while the differences might affect clinical-decision making for the others (LP vs VUS: 26/141; P vs VUS: 35/141; LP vs LB: 1/141; and 8 comparisons where the absence of a third label led to an additional difference). After excluding results for which this comparison was not possible (i.e., 8 comparisons of variants without a label), 91% (361/397) of the pairwise evaluations differed by at most one level (e.g. P vs. LP), and the remaining 9% (36/397) differed by at most two levels (e.g. P vs. VUS).

To evaluate potential causes of disagreement, various analyses were performed. Firstly, the distribution of the labels of the variants that were unanimously classified versus the ones where there was disagreement, were compared. The variants for which labelling was unanimous were proportionately positioned towards the extremes of the classification compared to the others (Fig. 2, 93% vs 46% received a P label, respectively). Secondly, the assigned labels were compared with the ones assigned by an automated decision algorithm that followed the decision table exactly. This led to the identification of 42 classification differences. After verification by the original evaluator, 11 labels were kept, while 31 others were changed. This led to the following distribution of labels: 301 P, 59 LP, 39 VUS, 1 LB, 1 B and 4 without a label (Suppl. File 5). The agreement on the classification increased to 69% (279/405) as 78 variants were unanimously categorized, while 45 and 12 variants received two or three different labels, respectively. Overall, out of 126 of 405 (31%) disagreements, 44% (55/126: P vs LP; 1/126: VUS vs B) were linked to the confidence of calls, while the differences might affect clinical-decision making for the others (P vs VUS: 38/126; LP vs VUS: 21/126; P vs LB: 2/126; P vs B: 1/126; and 8 comparisons where the absence of a third label led to an additional difference). Excluding the ones without a label, 89% (355/397 pairwise evaluations) of the pairwise comparisons differed at most by one level (e.g. P vs LP), by two levels for 10% (39/397, e.g. P vs VUS), and less than <1% of the comparisons by three (2/397, e.g. P vs LB) and four levels (1/397 e.g. P vs B), respectively. Thirdly, an evaluation of the individual criteria was performed. Twenty out of 23 criteria (all but BS3, BP3 and BP6) were used in the classification process, with a pairwise agreement of >80% for 14/23 and >70% for 19/23 criteria, respectively (Fig. 3A and B).

**Fig. 2.**
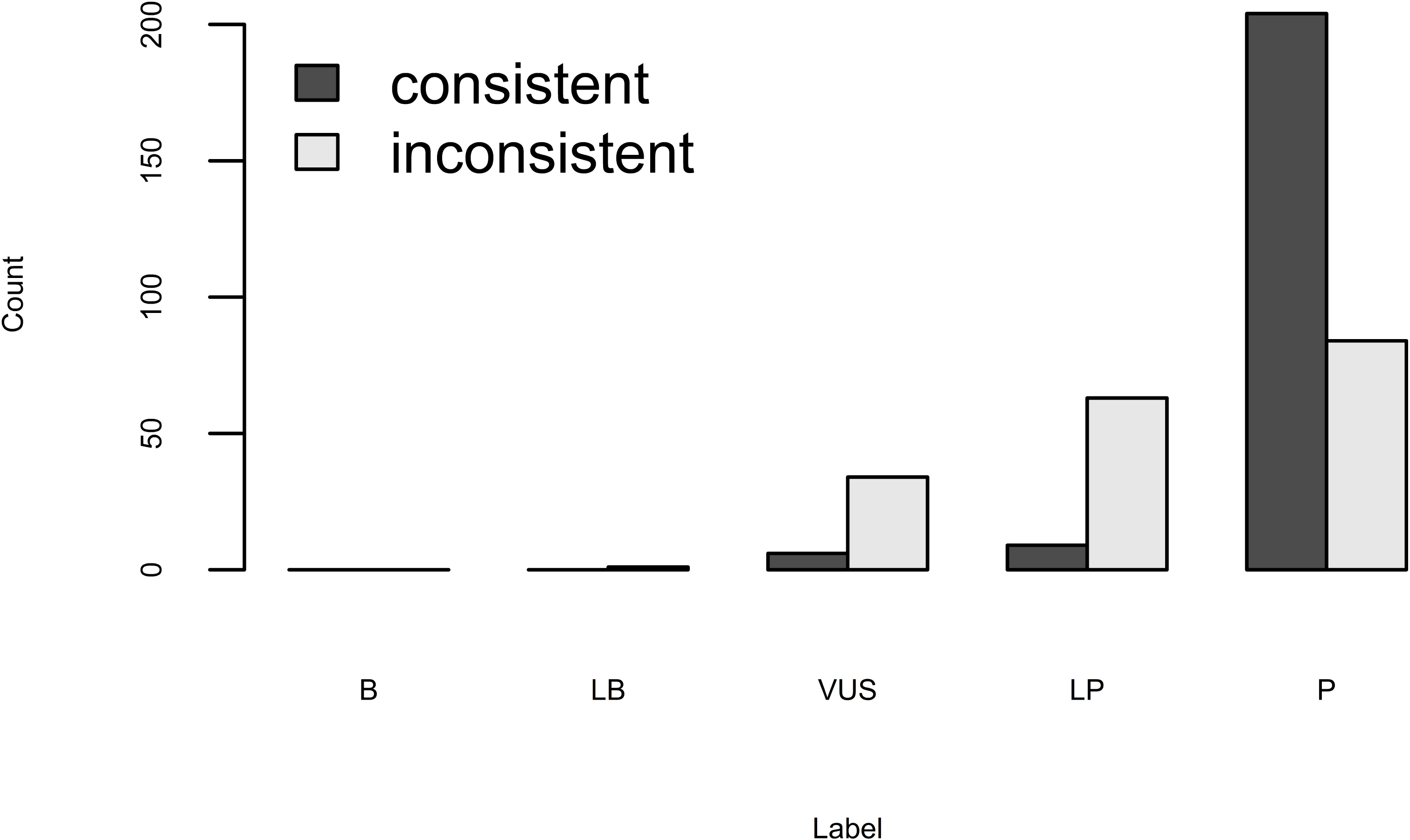
Distribution of the labels of the variants with a unanimous classification by all three reviewers versus the ones that received different labels. The label distribution was 3% VUS (6/219), 4% LP (9/219) and 93% P (204/219) for the variants that were unanimously classified, versus 1% LB (1/182), 19% VUS (34/182), 35% LP (63/182) and 46% P (84/182) for the variants where labels differed.

**Fig. 3.**
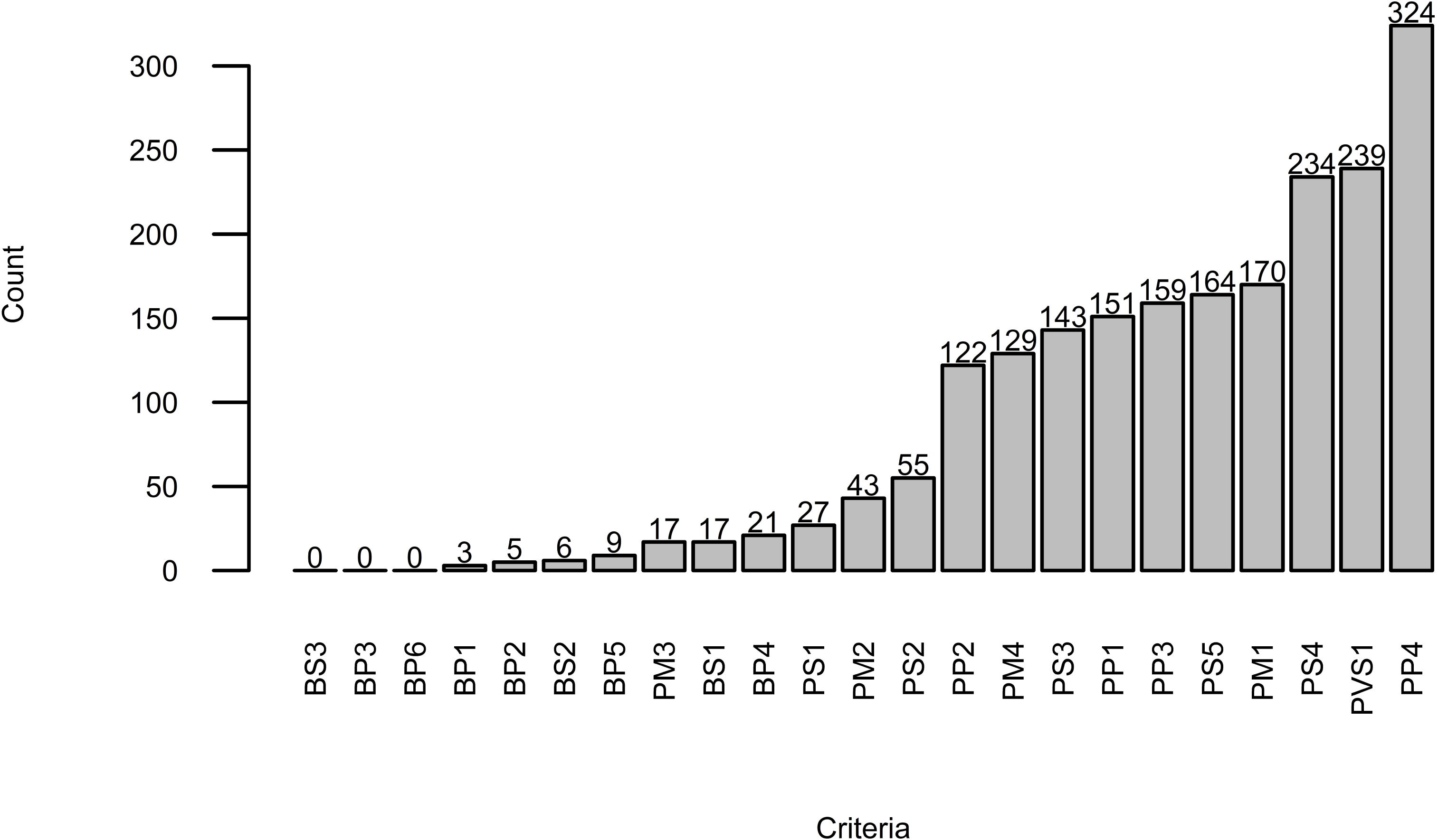
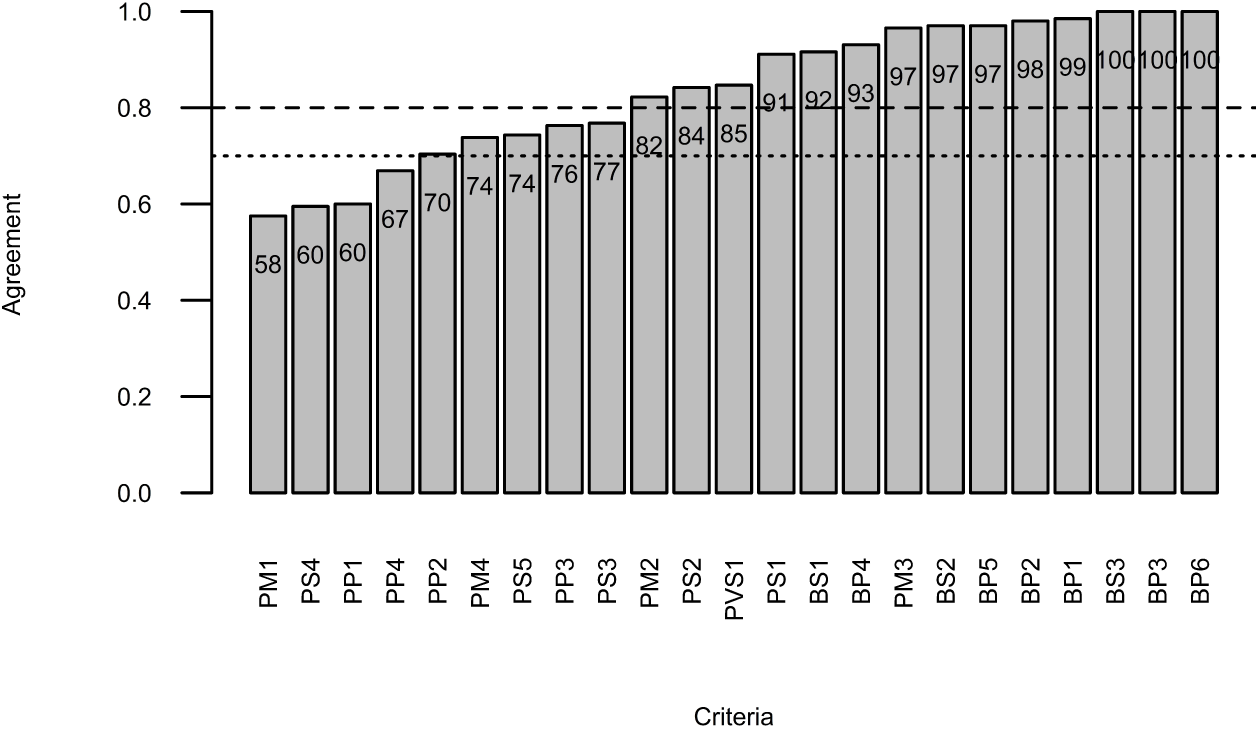
A. Graphical representation of the number of times individual criteria where used. Each bar mentions the number of times a specific criterion was used. B. Graphical representation of the interobserver agreement on the individual criteria (dashed line (- - -) = 80% agreement, dotted line (…) = 70% agreement). Each bar contains the observed agreement (%) for that specific criterion.

## Discussion

With the recent publication of AVCG, a new tool that aims to provide an objective and standardized methodology to classify animal genetic variants has become available (Boeykens et al., 2024a). As stated in the original paper and similar to the human ACMG/AMP guidelines, these guidelines are not final (and are not expected ever to become final); they should be continually reviewed and refined (Boeykens et al., 2024a; Richards et al., 2015). While a first step towards the evaluation of the reproducibility was included in the original publication, this was too limited to draw general conclusions: only 17 variants were evaluated, leading to 51 pairwise classifications (Boeykens et al., 2024a). Here, the reproducibility was thoroughly studied, with around ten times more variants, across three different species, and with a larger number of reviewers.

An important starting point in the process is determining whether variants actually fall in the scope and the extent to which this determination is reproducible across reviewers. To our knowledge, agreements on the scope have not been reported in the veterinary or human genetics literature before. Overall, with a pairwise agreement of 93%; decisions on scope were highly agreed on in the current study. The definition thus seems sufficiently clear to ensure a high reproducibility, as well as practically usable. Nevertheless, it was not perfect: e.g. 3 out of 4 *ABCB1* variants were considered outside the scope (Suppl. File 4 – 5). While reviewers thus tend to exclude pharmacogenomic variants, which is in line with the human ACMG/AMP guidelines, the clarity of the scope can likely be improved by explicitly mentioning whether pharmacogenomic variants can be classified with the current guidelines (Richards et al., 2015). Other variants were excluded for various reasons, e.g. because they were not considered to be a disease (e.g. some of the variants causing hypothrichia and roaning) or because a previous study using the guidelines already demonstrated that a variant did not cause disease (e.g. the *ALMS1* variant that was previously suggested to cause hypertrophic cardiomyopathy) (Boeykens et al., 2024b). In the latter case, the label was thus already available.

The most important part, i.e. label assignment, was quite consistent, with a 54% (73/135) unanimous agreement and 65% (264/405) pairwise agreement, which is in the same order of magnitude as the initial small study, where it was 74% (Boeykens et al., 2024a). In human literature, unanimous agreement based on the 2015 ACMG/AMP guidelines ranges from 34% to 83%, with a median of 54% (Table 1). Practically, this implies that the variant classification based on the published AVCG guidelines is identical to what is achieved with human variant classification. Quite intuitively, variants that received unanimous classifications resided proportionally more in the extreme categories, indicating more support from different criteria and/or more criteria with a heavier weight, compared to the ones that were less consistently classified (Fig. 2). In the more extreme categories, it is less likely that small interpretational differences will cause a shift, leading to an increased agreement overall.

**Table 1.**
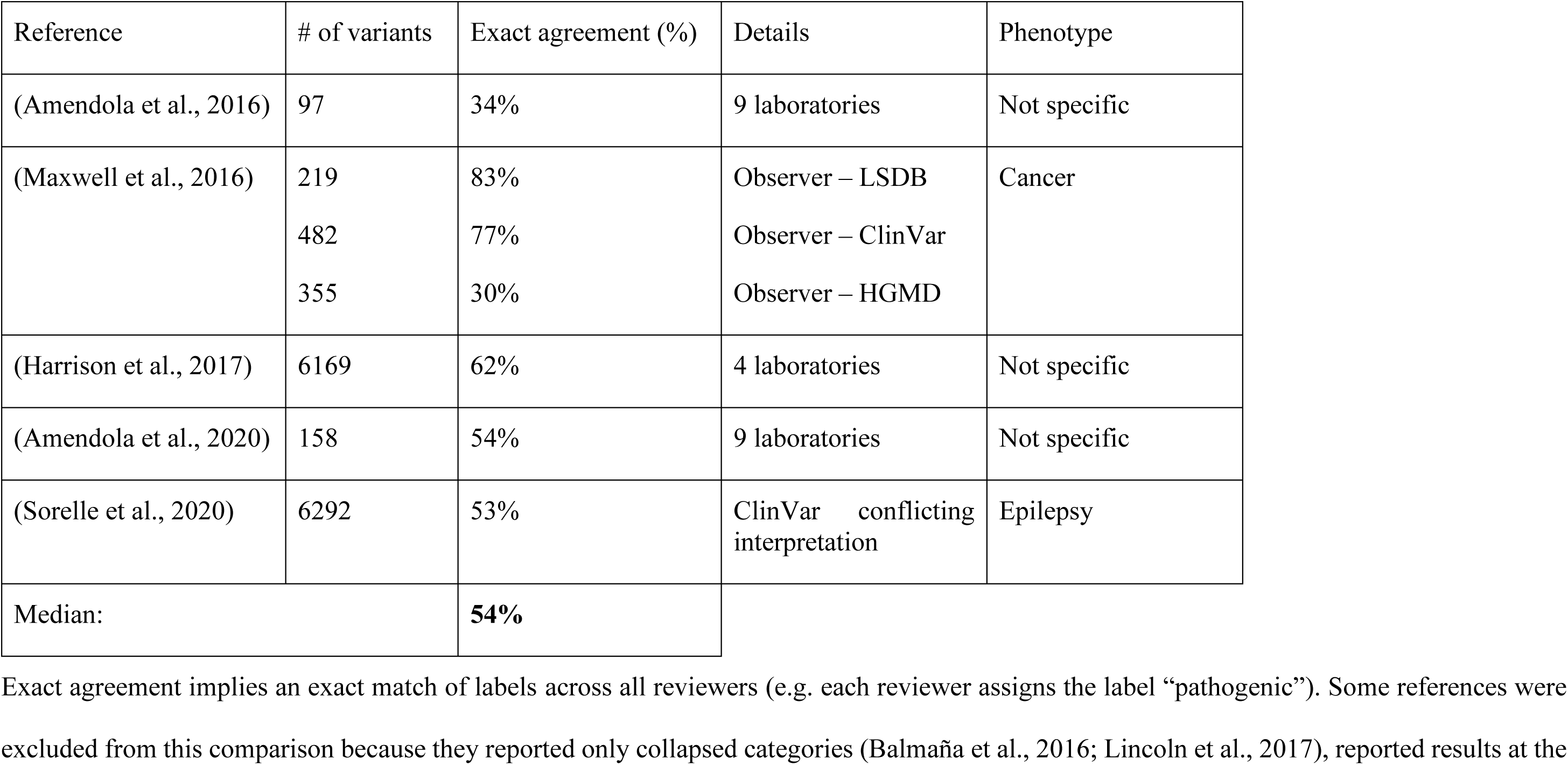
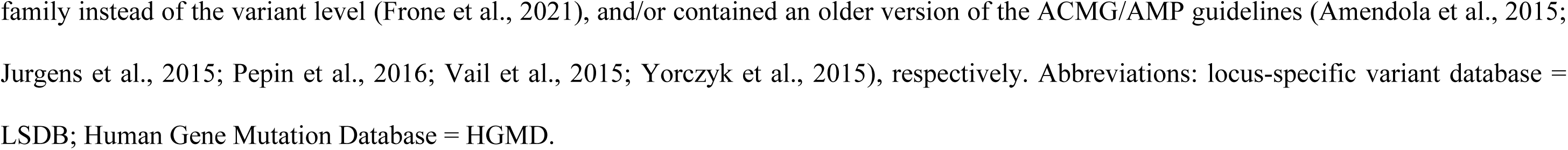
Chronological literature overview on the reproducibility of human genetic variant classification, since the 2015 version of the American College for Medical Genetics and Genomics and Association for Molecular Pathology guidelines (ACMG/AMP) (Richards et al., 2015).

| Reference | # of variants | Exact agreement (%) | Details | Phenotype |
| --- | --- | --- | --- | --- |
| (Amendola et al., 2016) | 97 | 34% | 9 laboratories | Not specific |
| (Maxwell et al., 2016) | 219 | 83% | Observer – LSDB | Cancer |
|  | 482 | 77% | Observer – ClinVar |  |
|  | 355 | 30% | Observer – HGMD |  |
| (Harrison et al., 2017) | 6169 | 62% | 4 laboratories | Not specific |
| (Amendola et al., 2020) | 158 | 54% | 9 laboratories | Not specific |
| (Sorelle et al., 2020) | 6292 | 53% | ClinVar conflicting interpretation | Epilepsy |
| Median: |  | **54%** |  |  |

Exact agreement implies an exact match of labels across all reviewers (e.g. each reviewer assigns the label “pathogenic”). Some references were excluded from this comparison because they reported only collapsed categories (Balmaña et al., 2016; Lincoln et al., 2017), reported results at the
family instead of the variant level (Frone et al., 2021), and/or contained an older version of the ACMG/AMP guidelines (Amendola et al., 2015; Jurgens et al., 2015; Pepin et al., 2016; Vail et al., 2015; Yorzcyk et al., 2015), respectively. Abbreviations: locus-specific variant database = LSDB; Human Gene Mutation Database = HGMD.

While an exact agreement of labels is the ultimate aim, some differences are more tolerated than others with respect to clinical decision-making. In more detail, P/LP differences or VUS/LB/B differences are less likely to influence clinical decision-making. Hence, almost all literature in human genetics predominantly focuses on what was defined here as clinically-relevant agreement (Amendola et al., 2020, 2016; Balmaña et al., 2016; Frone et al., 2021; Maxwell et al., 2016; Pepin et al., 2016; Sorelle et al., 2020). Disagreements between laboratories at this level can lead to e.g. some individuals in a family receiving the advice for a prophylactic treatment like a mastectomy (P/LP label), while others in the same family do not even receive the advice to perform additional screening and/or might not even be covered by insurance (VUS label) (Frone et al., 2021). In veterinary medicine, case reports and/or practical examples linked to these label differences have not been reported so far, probably due to the recent nature of the AVCG, however, with genetic counselling in veterinary medicine being more and more established (Adant et al., 2025), it is to be expected that case reports will be published in the near future. Overall, the agreement reported in (human genetics) literature increases quite substantially when it is assessed at this clinical level and similarly, the pairwise agreement increased here to 83% (335/405) (Amendola et al., 2020, 2016; Balmaña et al., 2016; Frone et al., 2021; Maxwell et al., 2016; Pepin et al., 2016; Sorelle et al., 2020). As a final comparison, the vast majority of the differences were at most a one-level difference (91% and 89% of the pairwise comparisons, respectively).

While agreement is thus as expected, the detailed analysis allowed the identification of several items that might further improve the results. Recommendations based on these findings are summarized in Table 2. The first one is that some labels were erroneously assigned based on tabulation errors. Also, in human literature, tabulation errors have been reported, albeit that the error rate here was slightly higher (74% (31/42)) versus 56% (9/16)) (Amendola et al., 2016). When the reviewers were asked to check these diverging labels, the ones that were changed, improved classification from 65% to 69%. While it is important to emphasize that reviewers can diverge from the decision table if they deem this necessary, the first recommendation based on this finding is to provide an automated label assignment in the files used to classify variants and this has already been implemented in the meantime (Table 2).

**Table 2.**
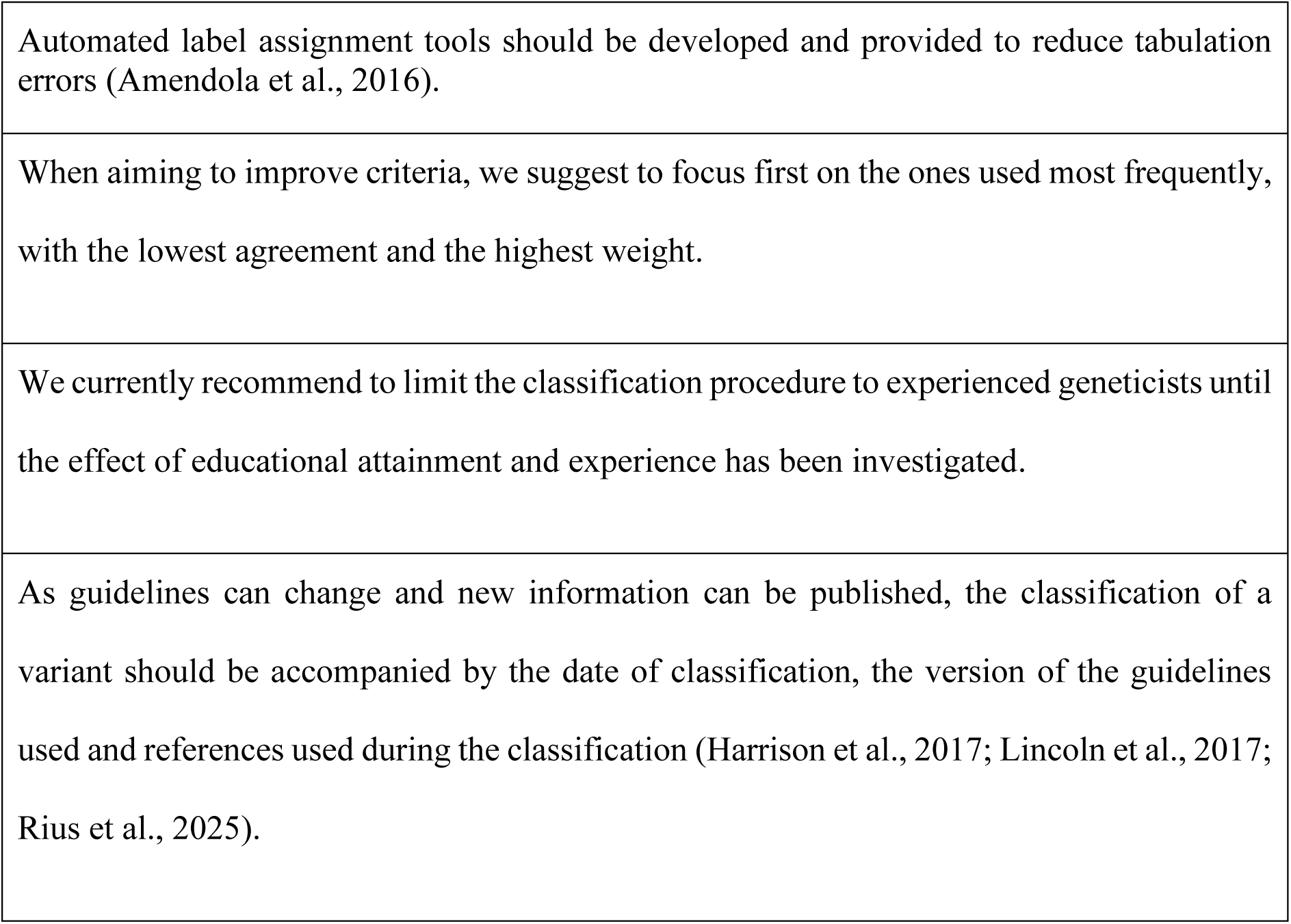
Recommendations to at least maintain and potentially further improve the current results. Additional references substantiating these recommendations are provided.

A detailed evaluation of the individual criteria demonstrated that some criteria (i.e. BP6, BP3 and BS3) were never used. A similar human study reporting the usage frequency had close to identical results as the two least used criteria corresponded to BP6 (called BP7 in ACMG/AMP) and BP3 (also called BP3 in ACMG/AMP) (Amendola et al., 2016). Furthermore, BS3 was also at the lower end of the frequency-of-use spectrum (Amendola et al., 2016) (Fig. 3A). In general, there seems to be a negative correlation between how many times a criterion is used and agreement, which is most intuitively explained for the criteria never used (Fig. 3B). If a criterion is never used, every reviewer agrees that it is consistently “not fulfilled”, implying agreement is automatically 100%. When criteria are used more frequently (only up to the other extreme of course), it becomes more likely to disagree, and this is also visible here (Fig. 3B). Nevertheless, there are some notable exceptions, e.g. PS4 is used second most, but has an agreement of well over 80 percent. On the other hand, PP4 is used most frequently, but scores at the lower end of the agreement. Practically, the current results enable a clear view of which criteria might need to be clarified further (Table 2).

To our knowledge, no studies have evaluated the effect of educational attainment or years of experience on genetic variant classification. Furthermore, most studies limited the details on the evaluator to reporting the laboratory that did the classification (Amendola et al., 2020, 2016, 2015; Balmaña et al., 2016; Frone et al., 2021; Harrison et al., 2017; Lincoln et al., 2017; Rehm et al., 2015; Rius et al., 2025; Sorelle et al., 2020; Vail et al., 2015; Yorczyk et al., 2015). Nevertheless, as the laboratories mentioned are described as being routinely involved in this process, we deem it likely that they were highly experienced. Similarly, the reviewers involved in this study all had at least a decade of professional experience in genetics. Taking the overall reproducibility into account, we hypothesize that the reproducibility will likely drop if a reviewer does not have ample experience. We currently recommend limiting the classification procedure to experienced geneticists until the effect of educational attainment and experience has been investigated (Table 2).

## Limitations

Variants were obtained from OMIA variant tables, and more specifically from the subset of “causal variants for single-gene diseases” (see materials and methods). These variants are derived from the published literature and have been deemed likely causal for a disease by one of three experienced curators. This was a deliberate choice, taking the ISAG AGTS initiative (i.e. to classify as many putative disease-causing variants as quickly as possible) and AVCG scope into account (i.e. disease-causing variants). However, this also implies that it is more likely that proportionately more disease-causing variants are included and far less variants from the LB/B spectrum. This is visible in the distribution of labels, with most of the variants being assigned a P/LP label. Furthermore, similar to another publication detailing criteria usage, not all criteria were used in the current study (Amendola et al., 2016). Future studies should focus on evaluating whether that might affect reproducibility.

## Conclusion

Overall, the current findings substantiate the reproducibility results of AVCG obtained in the first pilot study and are also in line with the results found in literature on the use of human ACMG/AMP guidelines. Furthermore, we identified several factors that might further improve reproducibility, including automated label calculation, and prioritization of criteria that might benefit most from additional clarification. Several of these recommendations have already been addressed or are currently being implemented. To ensure easy access and quick updating of labels whenever necessary, the labels will also be displayed in the OMIA variant tables.

## Author contributions

**Iris Casselman:** Investigation; Project Administration; Resources; Writing – Original Draft Preparation. **Carlotta Ferrari**: Investigation; Formal Analysis; Writing – Review & Editing. **Marie Abitbol:** Investigation; Formal Analysis; Writing – Review & Editing. **Danika Bannasch**: Investigation; Formal Analysis; Writing – Review & Editing. **Jerold Bell:** Investigation; Formal Analysis; Writing – Review & Editing. **Caroline Dufaure de Citres:** Investigation; Formal Analysis; Writing – Review & Editing. **Carrie J. Finno:** Investigation; Formal Analysis; Writing – Review & Editing. **Jessica J. Hayward:** Investigation; Formal Analysis; Writing – Review & Editing. **Jens Häggström:** Investigation; Formal Analysis; Writing – Review & Editing. **Jason T. Huff:** Investigation; Formal Analysis; Writing – Review & Editing. **Tosso Leeb:** Investigation; Formal Analysis; Writing – Review & Editing. **Ingrid Ljungvall:** Investigation; Formal Analysis; Writing – Review & Editing. **Maria Longeri:** Investigation; Formal Analysis; Writing – Review & Editing. **Leslie A. Lyons:** Investigation; Formal Analysis; Writing – Review & Editing. **Marcela Martinez:** Investigation; Formal Analysis; Writing – Review & Editing. **Cathryn Mellersh:** Investigation; Formal Analysis; Writing – Review & Editing. **Frank W. Nicholas:** Investigation; Formal Analysis; Writing – Review & Editing. **Åsa Ohlsson**: Investigation; Formal Analysis; Writing – Review & Editing. **Pascale Smets:** Investigation; Formal Analysis; Writing – Review & Editing. **Maria G. Strillacci:** Investigation; Formal Analysis; Writing – Review & Editing. **Imke Tammen**: Investigation; Formal Analysis; Writing – Review & Editing; OMIA database curator. **Frank G. van Steenbeek:** Investigation; Formal Analysis; Writing – Review & Editing. **Bart J.G. Broeckx:** Conceptualization; Data Curation; Formal Analysis; Software; Writing – Review & Editing; Supervision; Visualization; Funding Acquisition.

## Acknowledgements and funding

IC is partially funded by the Industrieel Onderzoeksfonds (IOF, F2024/IOF-ConcepTT/001) and Bijzonder Onderzoeksfonds Basic Research Funding (bof/baf/4y/2024/01/266) at Ghent University.

## Data Availability

The data that supports the findings of this study are available in the main text and supplementary material of this article.

## Ethics approval

No ethics approval was necessary for this study.

