## Supplementary material for "Reproducibility of the evaluation of genetic variant pathogenicity based on the animal variant classification guidelines": Suppl. File 1

Suppl. File 1. Criteria to support classification of pathogenic and benign variants in animals.

| Name | Criterion |
| --- | --- |
| PVS1 | Null variant (nonsense, frameshift, canonical ±1 or 2 splice-sites, initiation codon, single or multi-exon deletion) in a gene where LOF is a known mechanism of disease in the same or another species, if functionality of the gene is expected to be similar across species. |
| PS1 | Same amino acid change as a previously established pathogenic variant regardless of nucleotide change. |
| PS2 | *de novo* in a patient with the disease and unaffected parental samples tested negative. |
| PS3 | Well-established *in vitro* or *in vivo* functional studies supportive of a damaging effect on the gene or gene product. |
| PS4 | The prevalence of the variant in affected individuals is significantly increased compared with the prevalence in controls. |
| PS5 | Cosegregation with disease in multiple affected family members in a gene definitively known to cause the disease. |
| PM1 | Located in a mutational hot-spot and/or critical and well-established functional domain (e.g., active site of an enzyme) without benign variation across breeds and/or species. |
| PM2 | Novel missense change at an amino acid residue where a different missense change has been determined to be pathogenic in other individuals. |
| PM3 | For recessive disorders, detected in *trans* with a pathogenic variant. |
| PM4 | Protein length changes as a result of in-frame deletions/insertions in a non-repetitive region or stop-loss variants. |
| PP1 | Cross-species alignment shows the variant is conserved and other information across species (e.g., ClinVar data) states the variant is pathogenic. |
| PP2 | Missense variant in a gene that has a low rate of benign missense variation and in which missense variants are a common mechanism of disease. |
| PP3 | All computational evidence supports a deleterious effect on the gene or gene product (conservation, evolutionary, splicing impact, etc.). |
| PP4 | Patient’s phenotype or family history is highly specific for a disease with a single genetic etiology. |
| BS1 | Lack of segregation in affected members of a family. |
| BS2 | Observed in a healthy adult individual for a recessive (homozygous), dominant (heterozygous), or X-linked (hemizygous) disorder, with full penetrance expected at an early age. |
| BS3 | Well-established *in vitro* or *in vivo* functional studies show **no** damaging effect on protein function or splicing. |
| BP1 | Cross-species alignment shows the variant is not conserved and other information across species (e.g., ClinVar data) states the variant is benign. |
| BP2 | Observed in *trans* with a pathogenic variant for a fully penetrant dominant gene/disorder or observed in cis with a pathogenic variant in any inheritance pattern. |
| BP3 | In-frame deletions/insertions in a repetitive region without a known function. |
| BP4 | All computational evidence supports a benign effect on the gene or gene product (conservation, evolutionary, splicing impact, etc.). |
| BP5 | Variant found in a case with an alternate molecular basis for disease. |
| BP6 | A synonymous (silent) variant for which splicing prediction algorithms predict no impact to the splice consensus sequence nor the creation of a new splice site AND the nucleotide is not highly conserved. |

The name of each criterion designates whether it supports pathogenic or benign classification of a variant (P or B, respectively), followed by the weight of the support (VS = very strong; S = strong; M = moderate; P = supportive, respectively) and a number that represents the criterion. Abbreviations: LOF = loss-of-function
