## Supplementary material for "Reproducibility of the evaluation of genetic variant pathogenicity based on the animal variant classification guidelines": Suppl. File 2

Suppl. File 2. Decision rules to assign a genetic variant pathogenicity label.

| Step 1: always check branch A and branch B | |
| --- | --- |
| - Branch A: pathogenic variant | |
| Pathogenic (P) | (i) Very strong (PVS1) AND |
|  | - ≥1 strong (PS1 – PS5) OR |
|  | - ≥2 moderate (PM1 – PM4) OR |
|  | - 1 moderate (PM1 – PM4) AND 1 supporting (PP1 – PP4) OR |
|  | - ≥2 supporting (PP1– PP4) |
|  | (ii) ≥2 strong (PS1 – PS5) |
|  | (iii) 1 strong (PS1 – PS5) AND |
|  | - ≥3 moderate (PM1 – PM4) OR |
|  | - 2 moderate (PM1 – PM4) AND ≥2 supporting (PP1 – PP4) OR |
|  | - 1 moderate (PM1 – PM4) AND 4 supporting (PP1 – PP4) |
| Likely Pathogenic (LP) | (i) Very strong (PVS1) AND 1 moderate (PM1 – PM4) |
|  | (ii) 1 strong (PS1 – PS5) AND 1-2 moderate (PM1 – PM4) |
|  | (iii) 1 strong (PS1 – PS5) AND ≥2 supporting (PP1 – PP4) |
|  | (iv) 3 moderate (PM1 – PM4) |
|  | (v) 2 moderate (PM1 – PM4) AND ≥2 supporting (PP1 – PP4) |
|  | (vi) 1 moderate (PM1 – PM4) AND ≥4 supporting (PP1 – PP4) |
| - Branch B: benign variant | |
| Benign (B) | ≥2 strong (BS1 – BS3) |
| Likely Benign (LB) | (i) 1 strong (BS1 – BS3) AND 1 supporting (BP1 – BP6) |
|  | (ii) ≥2 supporting (BP1 – BP6) |
| Step 2: variants without a label or with a label from the pathogenic (P/LP) and benign (B/LB) branch: | |
| Variant of Uncertain Significance (VUS) | (i) not enough criteria met to assign label |
|  | (ii) labels from two conflicting branches assigned |

To assign a pathogenicity classification label, all criteria mentioned in Suppl. File 1, have to be evaluated. Only the criteria that are fulfilled, are used to assign a label. The number of criteria met in each weight category are counted. Once all criteria have been evaluated and a final count per weight category has been obtained, a variant pathogenicity label can be assigned based on the decision rules mentioned herein. This is a stepwise process in which first both branch A and branch B have to be considered. If no classification is assigned or if a classification from branch A and B is assigned, step two also has to be considered.
