## Supplementary material for "Reproducibility of the evaluation of genetic variant pathogenicity based on the animal variant classification guidelines": Suppl. File 3

Suppl. File 3. Description and consequences of each variant pathogenicity label in the five-category system.

| Label | Consequence |
| --- | --- |
| Pathogenic (P) | a healthcare provider can use molecular testing information in clinical decision-making, for breeding programs and/or screening. |
| Likely pathogenic (LP) | a health-care provider can use the molecular testing information in clinical decision-making when combined with other evidence of the disease in question, for breeding programs and/or screening. |
| Variant of uncertain significance (VUS) | not to be used in clinical decision-making, for breeding programs or screening. Efforts to resolve the classification of the variant as pathogenic or benign should be undertaken. |
| Likely benign (LB) | a healthcare provider can conclude that it is not the cause of the patient’s disorder when combined with other information. It should not be used for breeding programs and/or screening. |
| Benign (B) | a healthcare provider can conclude that it is not the cause of the patient’s disorder. It should not be used for breeding programs and/or screening. |

Each category is linked to specific recommendations in terms of clinical decision making, inclusion in breeding and screening programs. Between P and LP, mainly the strength of the evidence varies, while the practical differences are limited. Similarly, the practical differences between VUS, LB, B also have limited consequences. The most significant health and breeding decisions would occur when a variant switches between the pathogenic (P/LP) and the benign/uncertain classifications (VUS/LB/B), which can happen when new information becomes available or when label assignment differs between evaluators, hence the importance of a high inter-evaluator agreement on classification.
